# Mitochondrial metabolism in primary and metastatic human kidney cancers

**DOI:** 10.1101/2023.02.06.527285

**Authors:** Divya Bezwada, Nicholas P. Lesner, Bailey Brooks, Hieu S. Vu, Zheng Wu, Ling Cai, Stacy Kasitinon, Sherwin Kelekar, Feng Cai, Arin B. Aurora, McKenzie Patrick, Ashley Leach, Rashed Ghandour, Yuanyuan Zhang, Duyen Do, Jessica Sudderth, Dennis Dumesnil, Sara House, Tracy Rosales, Alan M. Poole, Yair Lotan, Solomon Woldu, Aditya Bagrodia, Xiaosong Meng, Jeffrey A. Cadeddu, Prashant Mishra, Ivan Pedrosa, Payal Kapur, Kevin D. Courtney, Craig R. Malloy, Vitaly Margulis, Ralph J. DeBerardinis

**Affiliations:** Children’s Medical Center Research Institute; Quantitative Biomedical Research Center; Department of Urology; Department of Pediatrics; Department of Radiology; Kidney Cancer Program; Department of Pathology; Department of Internal Medicine; Advanced Imaging Research Center; Howard Hughes Medical Institute University of Texas Southwestern Medical Center, Dallas, Texas 75390, USA.

**Author notes:** Corresponding author Lead Contact Information: Ralph J. DeBerardinis, M.D., Ph.D. 5323 Harry Hines Blvd. Dallas, TX 75390-8502.

## Abstract

Most kidney cancers display evidence of metabolic dysfunction^1–4^ but how this relates to cancer progression in humans is unknown. We used a multidisciplinary approach to infuse ^13^C-labeled nutrients during surgical tumour resection in over 70 patients with kidney cancer. Labeling from [U-^13^C]glucose varies across cancer subtypes, indicating that the kidney environment alone cannot account for all metabolic reprogramming in these tumours. Compared to the adjacent kidney, clear cell renal cell carcinomas (ccRCC) display suppressed labelling of tricarboxylic acid (TCA) cycle intermediates in vivo and in organotypic slices cultured ex vivo, indicating that suppressed labeling is tissue intrinsic. Infusions of [1,2-^13^C]acetate and [U-^13^C]glutamine in patients, coupled with respiratory flux of mitochondria isolated from kidney and tumour tissue, reveal primary defects in mitochondrial function in human ccRCC. However, ccRCC metastases unexpectedly have enhanced labeling of TCA cycle intermediates compared to primary ccRCCs, indicating a divergent metabolic program during ccRCC metastasis in patients. In mice, stimulating respiration in ccRCC cells is sufficient to promote metastatic colonization. Altogether, these findings indicate that metabolic properties evolve during human kidney cancer progression, and suggest that mitochondrial respiration may be limiting for ccRCC metastasis but not for ccRCC growth at the site of origin.

## Main Text

Mitochondrial alterations are a common feature of many kidney malignancies, and the mechanisms underlying mitochondrial anomalies vary amongst kidney cancer subtypes. In clear cell renal cell carcinoma (ccRCC), the most common form of kidney cancer, approximately 90% of tumours have biallelic inactivation of the von Hippel-Lindau (VHL) tumour suppressor. Loss of VHL leads to pseudohypoxic stabilization of HIFα subunits and chronic activation of HIF target genes^5,6^, many of which promote glycolysis and suppress glucose oxidation^7–9^. A subset of chromophobe RCCs (chRCCs) contain mutations in Complex I of the electron transport chain (ETC)^2^, and almost all oncocytomas accumulate defective mitochondria through somatic mutations in Complex I and impaired mitochondrial elimination programs^10–12^. Pathogenic mutations in metabolic enzymes like fumarate hydratase (FH) and succinate dehydrogenase (SDH) are initiating events in FH deficient renal cell cancer (RCC)^13^ and SDH-deficient RCC^14^, respectively. Although many tumours originating in the kidney display mitochondrial dysfunction, it is unclear how these mitochondrial anomalies impact nutrient metabolism in humans.

Intra-operative infusion of ^13^C-labeled nutrients and subsequent metabolite extraction and analysis of ^13^C labelling from surgically-resected samples can reveal metabolic differences between tumours and adjacent tissue and among different tumours from the same organ^15,16^. We previously reported suppressed contribution of glucose carbon to TCA cycle intermediates in five human ccRCCs, implying reduced glucose oxidation in these tumours. Here we studied why this phenotype occurs in human ccRCC, whether it characterizes kidney tumours more generally, and whether metabolic properties evolve during ccRCC progression to distant metastatic disease in patients. We infused patients with ^13^C-glucose, ^13^C-acetate and ^13^C-glutamine, capitalizing on the complementary views of the TCA cycle provided by these nutrients to produce a detailed analysis of mitochondrial metabolism in human cancer.

## Kidney cancers have variable glucose metabolism

Patients undergoing partial or radical nephrectomy for kidney cancer were administered a ^13^C-labeled nutrient through a peripheral intravenous line during surgery (Fig. 1A). After resection (typically 2-3 hours after the beginning of the infusion), tissue samples for metabolic analysis were chosen in consultation with the attending pathologist or pathology assistant. Using this approach, we studied 60 patients infused with [U-^13^C]glucose with various RCC subtypes, including 38 patients with ccRCC. The clinical features of these patients are shown in Extended Data Table 1. ccRCC tumours infused with [U-^13^C]glucose in this cohort exhibited a strong transcriptional correlation (R=0.864) with the TCGA KIRC data set reporting 446 ccRCC patients, indicating that the infused ccRCC tumours reported in this paper are reflective of ccRCC biology reported in earlier studies (Extended Data Fig 1A, Extended Data Fig 2, Extended Data Table 2). The labeling ratio of citrate m+2 (i.e. the fraction of citrate molecules containing two ^13^C nuclei) to pyruvate m+3 was lower in ccRCC samples compared to adjacent kidney, indicating a reduced contribution of glucose through the pyruvate dehydrogenase (PDH) reaction in ccRCC tumours (Fig. 1B,C; full isotopologue distributions from tissues and plasma are in Extended Data Table 3). Labeling in the tumours and renal cortex (hereafter, adjacent kidney) samples was variable, reflecting both inter-patient variability and regional labeling differences among samples from the same patient (Fig. 1D). When tumour labelling was compared to the adjacent kidney labelling in the same patient, only 1 of 28 patients with ccRCC displayed a statistically significant increase in the citrate m+2/pyruvate m+3 ratio in the tumour (Extended Data Fig. 1B). Ten patients did not have matched adjacent kidney tissue available for analysis. In addition to suppressed citrate m+2/pyruvate m+3 labeling ratios, total labeling of citrate and other TCA cycle intermediates (1-(m+0)) was also suppressed in ccRCCs (Fig. 1E); this metric incorporates all routes of label entry into the TCA cycle, and multiple turns of the cycle. Importantly, suppressed labeling of TCA cycle intermediates was not observed in all RCC subtypes (Fig. 1C, Extended Data Fig. S1C.), indicating that this property does not directly result from tumour residence in the kidney, and is not an artifact of the surgical procedure (Extended Data Table 1). To confirm that this is an intrinsic metabolic property of human ccRCC, we generated multiple viable agarose-embedded slices of kidney or ccRCC tissues from 6 patients and labeled them with [U-^13^C]glucose ex vivo in medium formulated to contain a nutrient content similar to human plasma^17^ (Fig. 1F). This revealed a similar degree of labeling suppression in citrate and malate as what was observed in the patients (Fig. 1G, Extended Data Fig. S1D).

**Figure 1:**
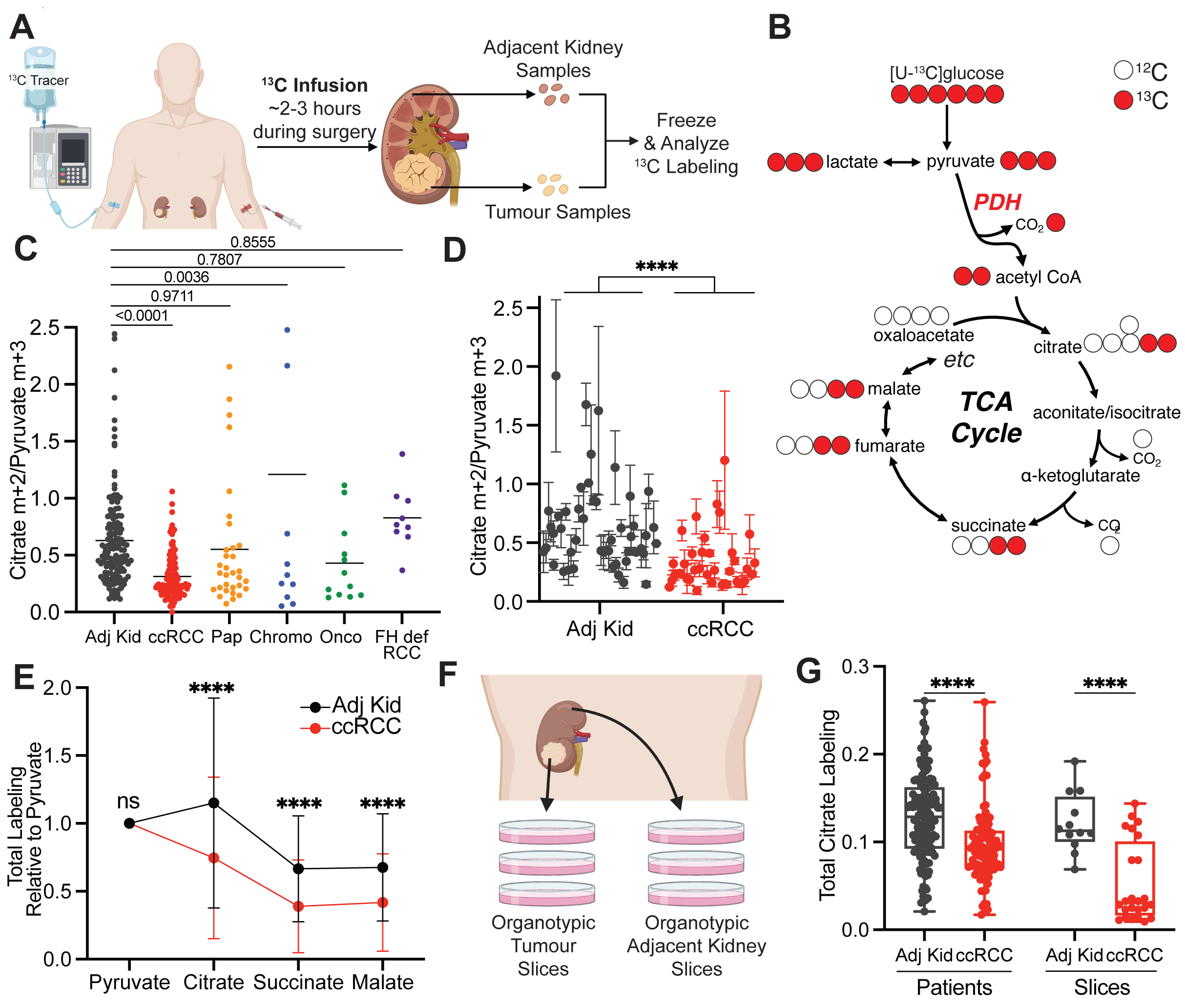
Glucose metabolism varies amongst kidney cancer subtypes. (A) Schematic of intraoperative infusions. **(B)** Schematic of isotopologue labeling in the tricarboxylic acid (TCA) cycle from [U-^13^C]glucose via pyruvate dehydrogenase (PDH). ^13^C carbons are indicated as red circles. **(C)** Citrate m+2/pyruvate m+3 ratio from patients infused with [U-^13^C]glucose. Each data point reflects an individual fragment of tissue. **(D)** Nested analysis of citrate m+2/pyruvate m+3 ratios separated by patient. Each data point represents a different patient. Error bars reflect the standard deviation from three fragments, tissue permitting, from the same patient. **(E)** Total isotopologue labeling (i.e. 1-(m+0)) of TCA cycle intermediates divided by total isotopologue labeling of pyruvate. **(F)** Schematic of organotypic patient tissue cultures. Tissue sections of ∼300 μM were placed on PTFE inserts in an incubator with 5% O2 for culture. **(G)** Total citrate labeling (1-(m+0)) from [U-^13^C]glucose in patients or tissue slices after 3 hours of labeling. All data represent mean ± standard deviation. Statistical significance was assessed using a one way analysis of variance (ANOVA) with a multiple comparison adjustment using Tukey’s methods (C), a nested t-test (D), or unpaired t-tests (E and G). \**P* < 0.05, \*\**P* < 0.01, \*\*\**P* < 0.001, \*\*\*\**P*<0.0001. Adj Kid = adjacent kidney, ccRCC = clear cell renal cell carcinoma, Pap = papillary renal cell carcinoma, Chromo = chromophone renal cell carcinoma, Onco = oncocytoma, FH def. RCC = FH deficient renal cell carcinoma. Fig 1A and 1F were created with biorender.com

**Figure 2:**
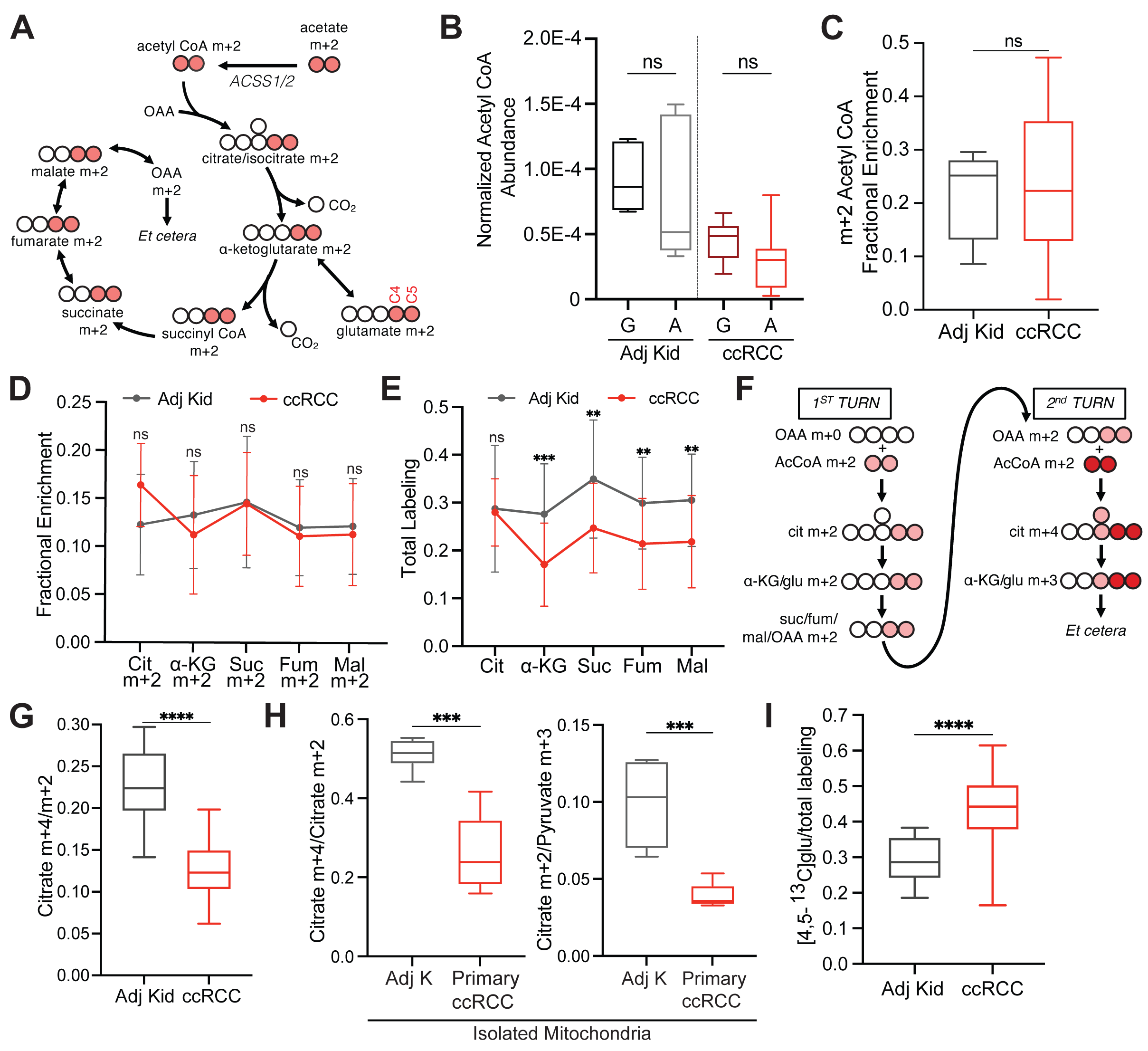
TCA cycle metabolism downstream of PDH is suppressed in ccRCCs. (A) Schematic of isotopologue labeling from [1,2-^13^C]acetate. **(B)** Total ion count (TIC)-normalized acetyl-CoA abundance after infusion with [U-^13^C]glucose (G) or [1,2-^13^C]acetate (A). **(C)** Enrichment of m+2 acetyl-CoA in the adjacent kidney versus ccRCC tumours. **(D)** m+2 isotopologues of TCA cycle intermediates from ccRCC patients infused with [1,2-^13^C]acetate. **(E)** Total labeling (1-(m+0)) of TCA cycle intermediates from ccRCC patients infused with [1,2-^13^C]acetate. **(F)** ^13^C labeling in the TCA cycle through two turns in the presence of [1,2-^13^C]acetyl-CoA. ^13^C from the first turn is in light red and ^13^C from the second turn is in dark red. **(G)** Citrate m+4/citrate m+2 ratios from the adjacent kidney and ccRCC tumours. **(H)** Citrate m+4/citrate m+2 and citrate m+2/pyruvate m+3 ratios from mitochondria isolated from the adjacent kidney or ccRCC tumours. **(I)** [4,5-^13^C]glutamate labeling as a fraction of total glutamate labeling after infusion with [1,2-^13^C]acetate. All data represent mean ± standard deviation, and whiskers of box and whisker plots represent minimum and maximum values. Statistical significance was assessed using unpaired two tailed parametric t-tests (B-E, G, H). ns *P>*0.05, \**P* < 0.05, \*\**P* < 0.01, \*\*\**P* < 0.001, \*\*\*\**P*<0.0001. Adj Kid = adjacent kidney, ccRCC = clear cell renal cell carcinoma.

## Acetate and glutamine supply the TCA cycle in ccRCC

We next infused 12 ccRCC patients with [1,2-^13^C]acetate (m+2), which can be converted to acetyl-CoA m+2 by acetyl-CoA synthetases (ACSS1/2, Fig. 2A). This tracer is useful for two reasons in this context. Unlike pyruvate, which can enter the TCA cycle through both acetyl-CoA and oxaloacetate (OAA) and produces complex labeling on even the first TCA cycle turn^18^, acetate only enters the TCA cycle through acetyl-CoA. This exclusively produces m+2 labeling in the first turn. Second, [1,2-^13^C]acetate transmits ^13^C to the TCA cycle independently of PDH, and so it is an informative complement to tracers like [U-^13^C]glucose that produce ^13^C-pyruvate, the substrate of PDH. The conditions we used to infuse [1,2-^13^C]acetate did not alter acetyl-CoA levels in tumours or adjacent kidneys and produced similar levels of acetyl-CoA labeling in both tissues (Fig. 2B,C, see Extended Data Table 4 for full isotopologue distributions). Fractional enrichments of m+2 TCA cycle intermediates in ccRCC tumours were also similar to adjacent kidney (Fig. 2D), indicating similar contributions to the TCA cycle under these infusion conditions. However, total labeling (1-(m+0)) of these metabolites revealed decreased labeling in tumours compared to kidney, consistent with reduced labeling beyond turn 1 of the TCA cycle (Fig 2E).

We then examined TCA cycle turnover in three complementary ways. First, the high enrichment in acetyl-CoA (average of 20-25%) allowed us to observe higher-order labeling in TCA cycle intermediates from subsequent rounds of incorporation of acetyl-CoA m+2 (Fig. 2F). The ratio of citrate m+4/m+2, a marker of ^13^C retention through two cycles, was reduced by about half in tumours relative to kidneys (Fig. 2G). Second, we examined labeling of TCA cycle intermediates in fresh mitochondria isolated from these resected tissues and cultured with [U-^13^C]pyruvate. Both the citrate m+2/pyruvate m+3 and citrate m+2/citrate m+4 ratios were decreased in the ccRCC mitochondria compared to kidney mitochondria (Fig. 2H), indicating that these metabolic properties are intrinsic to ccRCC mitochondria.

Third, we examined positional ^13^C labeling in glutamate, which exchanges with α-ketoglutarate and is classically used as a reporter of TCA cycle metabolism (Fig. 2A)^19–21^. Total glutamate labeling was lower in tumours compared to adjacent kidney (Extended Data Fig. 3A) but glutamate m+2 was similar (Extended Data Fig. 3B), mirroring the labeling pattern in TCA cycle intermediates. Strikingly, despite the similar m+2 fractional enrichment, the labeled ^13^C carbons were positioned differently in glutamate extracted from ccRCC tumours compared to the adjacent kidney. Using a tandem mass spectrometry method that reports isotopic position with high sensitivity^22^, we determined that [4,5-^13^C]glutamate, which appears in the first turn of the cycle (Fig. 2A), accounts for a much higher fraction of glutamate m+2 in tumours compared to adjacent kidney (Fig 2I). Therefore, most glutamate m+2 in ccRCC tumours comes from the first turn of the TCA cycle while labeling patterns requiring multiple turns of the TCA cycle are suppressed in ccRCC tumours.

**Figure 3:**
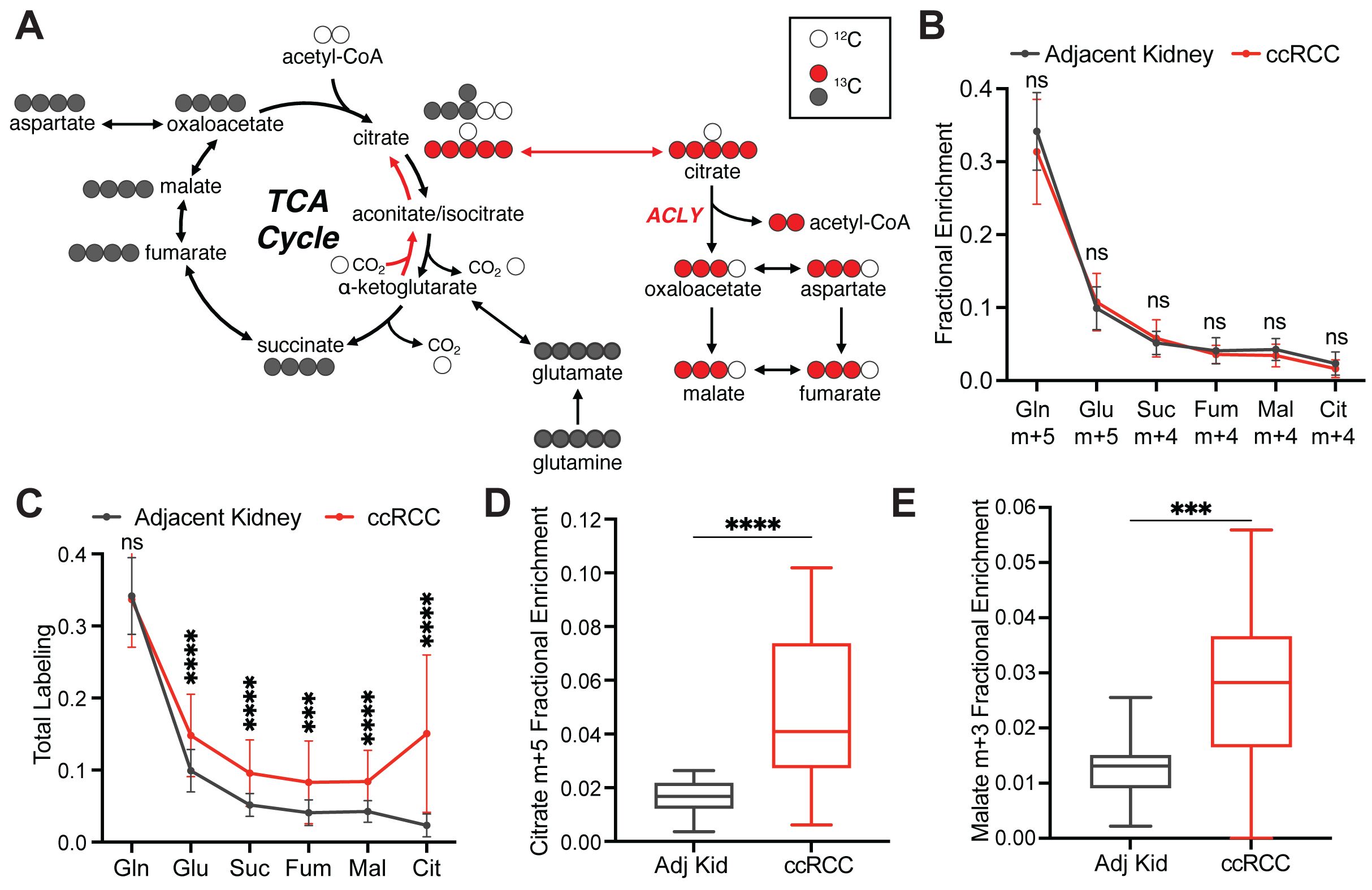
Glutamine contributes to the TCA cycle in ccRCC. **(A)** Schematic of isotopologue labeling from [U-^13^C]glutamine. Labeling from oxidative metabolism is indicated in grey and labeling from reductive metabolism is in red. **(B)** Isotopologues of TCA cycle intermediates from metabolism of [U-^13^C]glutamine through the first oxidative TCA cycle turn. **(C)** Total labeling (1-(m+0)) of TCA cycle intermediates from ccRCC patients infused with [U-^13^C]glutamine. **(D)** Fractional enrichment of m+5 citrate in the adjacent kidney and ccRCC tumours **(E)** Fractional enrichment of m+3 malate in the adjacent kidney and ccRCC tumours. Whiskers of box and whisker plots represent minimum and maximum values. Statistical significance was assessed using unpaired two tailed parametric t-tests (B-E). ns *P>*0.05, \**P* < 0.05, \*\**P* < 0.01, \*\*\**P* < 0.001, \*\*\*\**P*<0.0001. Adj Kid = adjacent kidney, ccRCC = clear cell renal cell carcinoma. Gln = glutamine, Glu = glutamate, Suc = succinate, Fum = fumarate, Mal = malate, Cit = citrate.

To assess the TCA cycle using a third tracer, we infused seven ccRCC patients with [U-^13^C]glutamine. Glutamine is the most abundant amino acid in the circulation, and its uptake in the tumour microenvironment is reported to be dominated by malignant cells^23^. Glutamine’s contributions to the TCA cycle involve conversion to alpha-ketoglutarate (α-KG) followed by either oxidation through α-KG dehydrogenase or reductive carboxylation by isocitrate dehydrogenase-1 or –2^24,25^. In cell culture, labeling through reductive metabolism is enhanced by processes that suppress pyruvate oxidation, including VHL loss, PDH suppression and mitochondrial defects^26–28^. Isotope labeling in citrate and other TCA cycle intermediates can discriminate which pathway is being utilized (Fig. 3A). The patient infusions produced the same glutamine m+5 enrichment in tumours and adjacent kidneys (30-35%, Fig. 3B). Labeling of glutamate m+5 and TCA cycle intermediates from the first turn of the oxidative TCA cycle (m+4) were also similar between tumour and kidney (Fig. 3B). However, total labeling (1-(m+0)) of these metabolites was higher in the tumours (Fig. 3C, see Extended Data Table 5 for full isotopologue distributions). The additional labeling in TCA cycle metabolites from the tumours involved enhanced contributions from the reductive pathway, as indicated by high citrate m+5 labeling in most fragments (Fig. 3D). This level of labeling far exceeded labeling in plasma citrate, indicating that it resulted from metabolism in the tumour (Extended Data Fig. 4A). The tumours also contained relatively high levels of malate m+3, indicating further metabolism along the reductive pathway (Fig. 3E). Therefore, glutamine is a carbon source in human ccRCC, and its metabolism results in oxidative and reductive labeling of TCA cycle intermediates.

**Figure 4:**
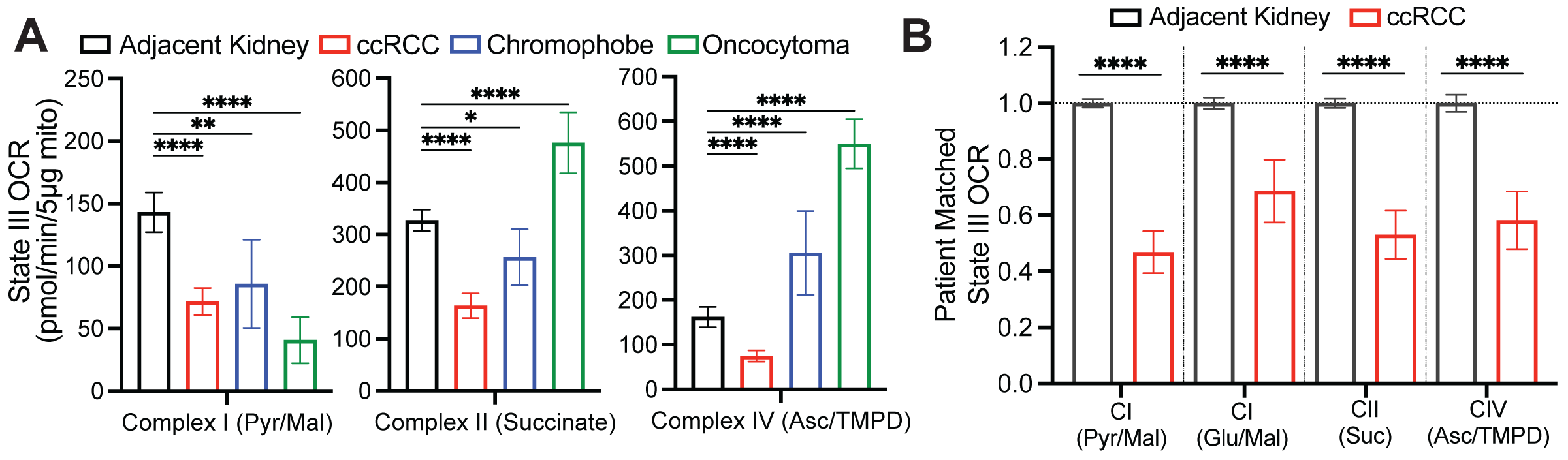
Respiration of mitochondria from primary human kidney cancers. **(A)** State III ADP-stimulated oxygen consumption rates (OCR) from mitochondria isolated from primary human tissues. Substrates used to stimulate respiration are indicated. **(B)** OCR from ccRCC mitochondria normalized to the patient matched adjacent kidney. Substrates used to stimulate respiration are indicated. Panels A and B represent mean ± 95% confidence intervals. Statistical significance was assessed using a one way analysis of variance (ANOVA) with a multiple comparison adjustment using Tukey’s methods (A) or unpaired two tailed parametric t-tests (B). ns *P>*0.05, \**P* < 0.05, \*\**P* < 0.01, \*\*\**P* < 0.001, \*\*\*\**P*<0.0001. Asc, ascorbate; TMPD = N,N,N,N-tetramethyl-p-phenylenediamine.

## Kidney cancers generally have low mitochondrial respiration

Suppressed labeling of TCA cycle intermediates from [U-^13^C]glucose and enhanced reductive labeling from [U-^13^C]glutamine are consistent with the effects of electron transport chain (ETC) dysfunction^26,29^. Multiple groups have reported decreases in mitochondrial DNA content^30–32^ and reduced expression of ETC components in RCC^33,34^. Transcriptional profiling from our cohort and the TCGA KIRC cohort both display reduced mRNA expression of ETC-related genes in ccRCC tumours relative to adjacent kidney, whereas many glycolytic genes are overexpressed in the tumours (Extended Data Fig. 1A). However, none of these analyses directly assessed coupled respiration in mitochondria from tumours and kidneys. We therefore measured oxygen consumption rates (OCR) of mitochondria immediately after harvesting them from fresh, surgically-resected kidney and tumour tissues. We used a differential centrifugation protocol to isolate mitochondria, then assessed ADP-stimulated (State III) and unstimulated (State IV) respiration. Mitochondria from both the kidney and ccRCC had normal respiratory control ratios (RCR, defined as State III/State IV respiration) when supplied with Complex I substrates, indicating that the preparation produced mitochondria with the expected ability to stimulate respiration upon addition of ADP^35^ (Extended Data Fig. 5A). However, absolute State III and State IV OCR was low in ccRCC compared to kidney mitochondria for complexes I, II, and IV (Fig. 4A, Extended Data Fig. 5B, Extended Data Fig. 5C). To account for day-to-day experimental variability in measuring respiration^36^, we also normalized OCR values from ccRCC mitochondria to patient-matched kidney mitochondria. From these 12 patients, the OCR from Complex I, II, and IV was always lower in mitochondria isolated from tumours (Fig. 4B).

**Figure 5:**
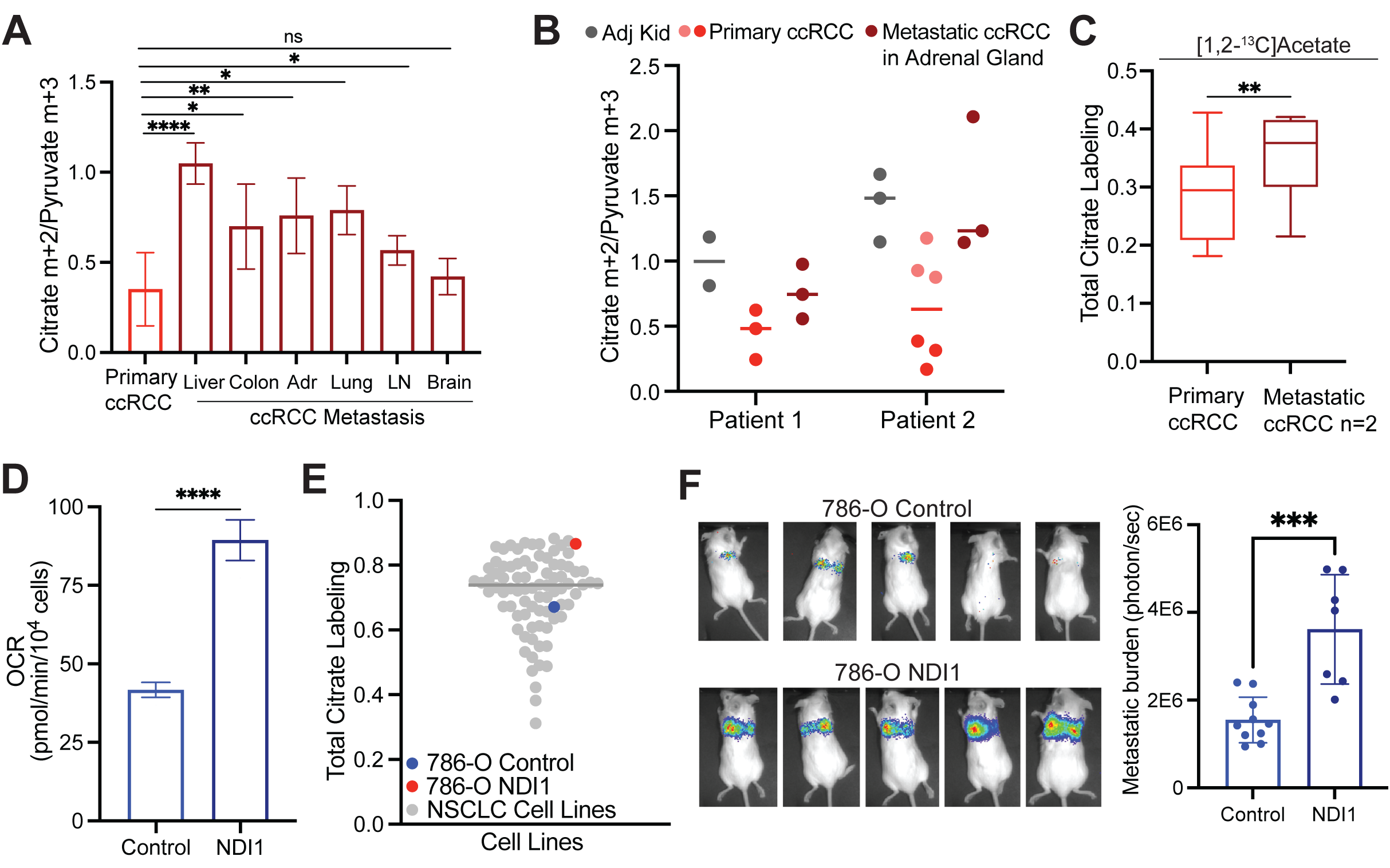
Metastatic ccRCCs utilize glucose differently than primary ccRCCs. **(A)** Citrate m+2/pyruvate m+3 ratio from patients infused with [U-^13^C]glucose. ccRCC metastases to different organ sites are indicated in dark red. **(B)** Citrate m+2/pyruvate m+3 ratio from two patients infused with [U-^13^C]glucose who had a primary ccRCC and synchronous metastasis to the adrenal gland removed during the same infusion. **(C)** Total citrate labeling (i.e 1-(m+0)) from patients infused with [1,2-^13^C]acetate. **(D)** OCR from 786-O control cells and 786-O cells expressing NDI1. **(E)** Total citrate labeling (i.e. 1-(m+0)) from cells cultured with [U-^13^C]glucose for 6 hours in RPMI with 5% dialyzed FBS. Labelling from non-small cell lung cancer (NSCLC) cell lines was previously published^49^. **(F)** Representative mice 4 weeks after tail vein injection of control and NDI1-expressing 786-O cells. Bioluminescence is quantified on the right. All data represent mean ± standard deviation. Statistical significance was assessed using unpaired two tailed parametric t-tests. Adr, adrenal gland; LN, lymph node. ns *P>*0.05, \**P* < 0.05, \*\**P* < 0.01, \*\*\**P* < 0.001, \*\*\*\**P*<0.0001.

Mitochondria from other RCC subtypes displayed low state III respiration at Complex I, but variable activities of other ETC components (Fig. 4A). Chromophobe tumours and oncocytomas contain mutations in genes encoding Complex I subunits, and accordingly both had low Complex I activity relative to adjacent kidney (Fig 4A). The RCRs of these mitochondria were also low when provided with Complex I substrates (Extended Data Fig. 5D). However, absolute state III OCRs for Complex II and IV were variable, and in mitochondria from oncocytomas, they exceeded rates from kidney mitochondria. Therefore, oncocytomas and chromophobe tumours display the expected defects in Complex I, with relative preservation of some other ETC components.

## Metastatic ccRCC tumours have increased TCA cycle labeling

Kidney cancer patients with early stage disease have a 5-year survival rate close to 95%. As in many cancers, ccRCC patients with distant metastases fare much worse, with 5-year survival rates under 15%^37^. How emergent metabolic properties support metastasis is a subject of intense investigation^38–46^ Most human studies describing metabolic alterations during metastasis are based on transcriptional data rather than direct assessment of metabolism in tumours^47,48^. Primary and metastatic human tumours have not been systematically compared using ^13^C infusions.

To directly examine metabolism in metastatic ccRCC, [U-^13^C]glucose was infused in 10 patients undergoing metastasectomy. Metastatic tumours in 9 of these 10 patients had higher citrate m+2/pyruvate m+3 ratios than the average citrate m+2/pyruvate m+3 ratio from primary ccRCCs from the kidney (Fig. 5A). Two patients with a primary ccRCC and a synchronous adrenal metastasis underwent concurrent nephrectomy and adrenalectomy, allowing both lesions to be sampled during the same infusion. Compared to the primary lesion, the metastatic adrenal tumours trended towards higher citrate m+2/pyruvate m+3 ratios compared to the primary tumour (Fig. 5B). In patient 2, two different regions of the primary tumour were sampled, with one region having a reduced citrate m+2/pyruvate m+3 ratio relative to the other region; both these regions had somewhat lower ratios than the metastasis (Fig 5B). Two patients with metastatic tumours were infused with [1,2-^13^C]acetate, and these tumours also displayed elevated citrate labeling compared to their matched primary ccRCCs (Fig. 5C). These data indicate that both glucose and acetate make larger contributions to the TCA cycle in metastatic than primary ccRCC, even in the same patients. To test for mechanisms to explain this observation, we performed RNA sequencing on non-tumour bearing kidney, primary ccRCC, and metastatic ccRCC nodules from seven patients. RNA sequencing did not show consistent alterations in transcripts associated with mitochondrial function or mtDNA content between primary and metastatic tumours (Extended Data Fig. 6A, Extended Data Fig 6B). The small size of these metastatic tumours precluded direct analysis of mitochondrial respiration.

The low apparent oxidative metabolism in primary ccRCCs provided an opportunity to test whether stimulating respiration would enhance metastatic spread. We expressed the yeast mitochondrial NADH dehydrogenase NDI1 in *VHL*-deficient 786-O ccRCC cancer cells. NDI1 oxidizes NADH to NAD+ and transfers electrons to the Coenzyme Q (CoQ) pool, essentially replacing the functions of mammalian Complex I. NDI1 expression enhanced respiration, rendered O2 consumption insensitive to Complex I inhibition (Fig. 5D, Extended Data Fig 6C) and increased labeling of TCA cycle intermediates from [U-^13^C]glucose (Extended Data Fig. 6D). We compared labeling of citrate from [U-^13^C]glucose for 6 hours in 786-O cells with and without NDI-1 to a panel of 81 non-small cell lung cancer cell lines that had been subjected to the same isotope labeling procedure^49^. Parental 786-O cells had below average citrate labeling, but NDI1-expressing cells had among the top 5% of citrate labeling (Fig. 5E). To assess the impact of NDI1 on metastatic colonization, the cells were engineered to express dsRed-luciferase and transplanted into immune compromised mice via the tail vein. Bioluminescence imaging revealed that NDI1 expression induced a large increase in lung colonization and growth as compared to 786-O cells expressing the empty vector (Fig 5F). Therefore, human metastatic ccRCCs display evidence of enhanced mitochondrial metabolism in patients, and ccRCC cells engineered to have increased oxidative phosphorylation display increased metastatic colonization in mice.

## Discussion

Two key points distinguish this work from prior studies on isotope tracing in human cancers. First, whereas earlier studies in other types of human cancer emphasized substantial TCA cycle labeling from [U-^13^C]glucose^15,50–52^, ccRCCs generally have low labeling relative to the adjacent kidney. We provide evidence that this is an intrinsic characteristic of ccRCC. Not all types of tumours growing in the kidney display suppressed glucose contribution to the TCA cycle, and importantly, we observe low glucose contributions to the TCA cycle in cultured slices of ccRCC tissue. Although these in vivo isotope infusions do not report quantitative fluxes, data from three different nutrient tracers ([U-^13^C]glucose, [1,2-^13^C]acetate and [U-^13^C]glutamine) are consistent with primary ccRCCs having suppressed TCA cycle turnover relative to adjacent kidney. While PDH suppression is a well-known effect of HIF-1α activation, we also report dysfunction of multiple ETC components manifesting as reduced mitochondrial respiration. This finding may be related to suppressed mtDNA copy number in ccRCC^30^, and it predicts that activating PDH would not be sufficient to normalize oxidative metabolism in ccRCC. Second, we report higher contributions of glucose to the TCA cycle in metastatic ccRCC compared to primary ccRCC. This was observed in both synchronous and asynchronous metastases, in multiple metastatic sites, and it implies an evolution or selection of mitochondrial function during ccRCC metastasis in patients.

Evidence from mice indicates that the TCA cycle and oxidative phosphorylation may promote multiple aspects of cancer progression, including metastasis. Quantitative measurement of TCA cycle flux in orthotopic models of breast cancer reported a large increase in flux after metastasis to the lung^52^. In melanoma, the formation and growth of brain metastases in mice is suppressed by inhibiting ETC Complex I^53^. The mechanisms underlying these effects are unknown, both in the mouse models and in our work in ccRCC patients. It is unclear whether tumour cells with variable mitochondrial function at the primary site activate mitochondrial metabolism during metastasis, or whether metastasis selects for a pre-existing population of cells with high mitochondrial metabolism. We have not pinpointed when and where in the metastatic cascade oxidative phosphorylation exerts its benefits for metastasis. However, our finding that NDI1 promotes tumour burden in the lung after tail vein injection suggests that part of the benefit occurs after escape from the primary tumour.

Other studies that did not focus explicitly on metastasis have also reported the differential importance of oxidative phosphorylation in advanced cancers. In a mouse model of pancreatic ductal adenocarcinoma, oxidative phosphorylation underlies relapse and outgrowth after genetic ablation of the oncogenic driver. In these mice, relapse is suppressed and survival is enhanced by inhibiting the ETC^54^. In acute myelogenous leukemia, human-derived mouse models with robust oxidative phosphorylation display resistance to cytotoxic chemotherapy, and this resistance is reversed by inhibiting mitochondrial function^55^. In patient-derived B-progenitor acute lymphoblastic leukemia models, clones destined to relapse have gene expression signatures of mitochondrial metabolism and higher mitochondrial mass than clones that do not relapse^56^. These findings suggest that oxidative phosphorylation and other aspects of mitochondrial function underlie a program of enhanced fitness that allows tumour cells to survive a variety of stresses relevant to cancer progression, including stresses related to metastasis.

Efforts to suppress cancer progression by targeting mitochondrial metabolism will benefit from understanding the basis of the relationship between the mitochondria and metastasis. It is unclear why ccRCC metastases in patients bear hallmarks of enhanced mitochondrial function, because neither the mtDNA content nor the levels of transcripts related to oxidative phosphorylation differed between primary and metastatic ccRCC in our cohort. Perhaps the most interesting and important challenge arising from this work is to determine which metabolic effects of mitochondrial function support metastasis. The ETC supports efficient ATP production from nutrient oxidation, and this may be essential to survive the reduced nutrient uptake that accompanies loss of anchorage^38,57^. But the ETC also supports the maintenance of a favorable redox balance, ubiquinol oxidation, and production of anabolic precursors, all of which support tumour growth in various contexts^58–61^. Potent, systemic blockade of the ETC in patients results in dose-limiting toxicities^62^, but it may be possible to widen the therapeutic window by tailoring therapies to selectively target the most relevant aspects of mitochondrial function.

## Acknowledgements

We are grateful to Gerardo Guevara for his efforts on this project. We thank Aron Jaffe for critiquing the manuscript. This article is subject to HHMI’s Open Access to Publications policy. HHMI lab heads have previously granted a nonexclusive CC BY 4.0 license to the public and a sublicensable license to HHMI in their research articles. Pursuant to those licenses, the author-accepted manuscript of this article can be made freely available under a CC BY4.0 license immediately upon publication. R.J.D. is supported by the Howard Hughes Medical Institute Investigator Program, grants from the N.I.H. (R35CA220449, P50CA196516) and Cancer Prevention and Research Institute of Texas (RP180778), and by the Moody Foundation (Robert L. Moody, Sr. Faculty Scholar Award). D.B. was supported by grants from the N.I.H (F31CA239330, T32GM008203, TL1TR001104). N.P.L was supported by grants from the N.I.H (F31DK122676). K.D.C is supported by P50CA196516. I.P was supported by grants from the N.I.H. (R01CA154475, U01CA207091, P50CA196516). Resources from P30CA142543 and P41EB15908 were used to support this study. The content of this manuscript is solely the responsibility of the authors and does not necessarily represent the official views of the NIH.

## Contributions

Conceptualization: D.B., R.J.D. Writing of manuscript: D.B., R.J.D. Project Supervision: V.M., R.J.D. Investigation: D.B., N.P.L., B.B., H.S.V., Z.W., L.C., S.K., S.K., F.C., A.A., M.P., A.L., R.G., Y.Z., D.D., J.S., D.D., S.S., T.R., A.M.P., Y.L., S.W., A.B., X.M., J.A.C., I.P., P.K., K.D.C., C.R.M., V.M., and R.J.D. Funding: R.J.D. All authors reviewed the manuscript.

## Methods

### Patient Infusions

Patients 18 years or older with radiographic evidence of known or probable kidney cancer requiring surgical biopsy or excision were recruited to an IRB-approved study and informed consent was obtained. Patients receiving [U-^13^C]glucose were enrolled on protocol STU062010-157 or STU2019-1061 and infused at the following rate: 8 gram bolus of [U-^13^C]glucose administered over 10 minutes, followed by a continuous infusion of [U-^13^C]glucose at either 4 or 8 grams/hour. Patients receiving [1,2-^13^C]acetate were enrolled on protocol STU2019-1061 and infused at the following rate: bolus of 3 mg [1,2-^13^C]acetate/kg/minute for 5 minutes, followed by a continuous infusion of [1,2-^13^C]acetate at 1.5 mg/kg/minute. Patients receiving [U-^13^C]glutamine were enrolled on protocol STU2019-1061 and were infused at the following rate: primer dose for 5 minutes at a rate of 0.6 mg/kg/minute, followed by a continuous infusion of 5.0 μmol/kg/minute (0.73 mg/kg/minute). Uncontrolled or poorly controlled diabetes and pregnancy were exclusion criteria for the study. Demographic, clinical and pathological details are summarized in Extended Data Table 1.

### Animal Studies

All procedures were approved by UT Southwestern Medical Center’s Animal Care and Use Committee in accordance with the Guide for the Care and Use of Laboratory Animals.

### Cell Lines

Cell lines were purchased from ATCC and confirmed to be mycoplasma free using the (Bulldog Bio, Cat. No. 2523348). Cells were maintained in RPMI supplemented with 10% fetal bovine serum or 10% dialyzed human serum and cultured at 37°C in 5% CO2 and 95% air, unless otherwise noted.

### High-resolution Mass Spectrometry (QTOF)

Data acquisition from isolated mitochondria, patient plasma, and patient tissues was performed by reverse-phase chromatography on a 1290 UHPLC liquid chromatography (LC) system interfaced to a high-resolution mass spectrometry (HRMS) 6550 iFunnel Q-TOF mass spectrometer (MS) (Agilent Technologies, CA). The MS was operated in both positive and negative (ESI+ and ESI-) modes. Analytes were separated on an Acquity UPLC® HSS T3 column (1.8 μm, 2.1 x 150 mm, Waters, MA). The column was kept at room temperature. Mobile phase A composition was 0.1% formic acid in water and mobile phase B composition was 0.1% formic acid in 100% ACN. The LC gradient was 0 min: 1% B; 5 min: 5% B; 15 min: 99%; 23 min: 99%; 24 min: 1%; 25 min: 1%. The flow rate was 250 μL min^-1^. The sample injection volume was 5 μL.

ESI source conditions were set as follows: dry gas temperature 225 °C and flow 18 L min^-1^, fragmentor voltage 175 V, sheath gas temperature 350 °C and flow 12 L min^-1^, nozzle voltage 500 V, and capillary voltage +3500 V in positive mode and −3500 V in negative. The instrument was set to acquire over the full *m/z* range of 40–1700 in both modes, with the MS acquisition rate of 1 spectrum s^-1^ in profile format. Raw data files (.d) were processed using Profinder B.08.00 SP3 software (Agilent Technologies, CA) with an in-house database containing retention time and accurate mass information on 600 standards from Mass Spectrometry Metabolite Library (IROA Technologies, MA) which was created under the same analysis conditions. The in-house database matching parameters were: mass tolerance 10 ppm; retention time tolerance 0.5 min. Peak integration result was manually curated in Profinder for improved consistency and exported as a spreadsheet (.csv).

### High-resolution Mass Spectrometry (Orbitrap)

[1,2-^13^C]acetate patient tissue samples were analyzed using an Orbitrap Fusion Lumos 1M Tribrid Mass Spectrometer. HILIC chromatographic separation of metabolites was achieved using a Millipore ZIC-pHILIC column (5 μm, 2.1 × 150 mm) with a binary solvent system of 10 mM ammonium acetate in water, pH 9.8 (solvent A) and acetonitrile (solvent B) with a constant flow rate of 0.25 ml min−1. For gradient separation, the column was equilibrated with 90% solvent B. After injection, the gradient proceeded as follows: 0–15 min linear ramp from 90% B to 30% B; 15–18 min isocratic flow of 30% B; 18–19 min linear ramp from 30% B to 90% B; 19–27 column regeneration with isocratic flow of 90% B. HRMS data were acquired with two separate acquisition methods. Individual samples were acquired with an HRMS full scan (precursor ion only) method switching between positive and negative polarities. For data-dependent, high-resolution tandem mass spectrometry (ddHRMS/MS) methods, precursor ion scans were acquired at a resolving power of 120,000 full width at half-maximum (FWHM) with a mass range of either 50-750 or 70-1,050 Da. The AGC target value was set to 1 × 10^6^ with a maximum injection time of 100 ms. Pooled samples were generated from an equal mixture of all individual samples and analyzed using individual positive- and negative-polarity spectrometry ddHRMS/MS acquisition methods for high-confidence metabolite ID. Product ion spectra were acquired at a resolving power of 15,000 FWHM without a fixed mass range. The AGC target value was set to 2 × 10^5^ with a maximum injection time of 150 ms. Data-dependent parameters were set to acquire the top 10 ions with a dynamic exclusion of 30 s and a mass tolerance of 5 ppm. Isotope exclusion was turned on and a normalized collision energy value of 30 was used or a stepped normalized collision energy applied with values of 30, 50 and 70. Settings remained the same in both polarities. Metabolite identities were confirmed in three ways: (1) precursor ion m/z was matched within 5 ppm of theoretical mass predicted by the chemical formula; (2) fragment ion spectra were matched within a 5 ppm tolerance to known metabolite fragments; and (3) the retention time of metabolites was within 5% of the retention time of a purified standard run with the same chromatographic method. Metabolites were relatively quantitated by integrating the chromatographic peak area of the precursor ion searched within a 5 ppm tolerance.

Acetyl-CoA fractional enrichment was determined with a selected ion monitoring (SIM) scan event on an Orbitrap Fusion Lumos 1M Tribrid Mass Spectrometer. The SIM scan event targeted the theoretical mass for the positive ion of acetyl-CoA in positive ionization mode (m/z 810.1330) with a 4.5 dalton window. Data was collected with a resolving power of 60,000 FWHM with an AGC target of 4E5 ions. To calculate fractional enrichment of M+2 acetyl-CoA, the SIM scan integrated the M+0, M+1 and M+2 peaks and the full scan data to integrate the remaining naturally abundant isotopes. Isotope enrichment was corrected for natural abundance.

### Isotopomer Analysis

Samples were analyzed on an AB Sciex 6500 QTRAP liquid chromatography/mass spectrometer (Applied Biosystems SCIEX) equipped with a vacuum degasser, quaternary pump, autosampler, thermostatted column compartment and triple quadrupole/ion trap mass spectrometer with electrospray ionization interface, and controlled by AB Sciex Analyst 1.6.1 Software. SeQuant® ZIC®-pHILIC 5µm polymer (150mm×2.1mm) columns were used for separation. Solvents for the mobile phase were 10 mM ammonium acetate aqueous (pH 9.8 adjusted with NH3.H2O (A) and pure acetonitrile (B). The gradient elution was: 0–20 min, linear gradient 90-65% B, 20– 23 min, linear gradient 65-30% B, 23-28 min, 30% B, and 28–30 min, linear gradient 30-90% B then reconditioning the column with 90% B for 5 min. The flow-rate was 0.2 ml/min and the column was operated at 40°C.

### Gas Chromatography-Mass Spectrometry (GC/MS)

GC/MS was used to analyze infused patient tissue and plasma samples as well as tracing assays in cell lines and slice cultures. Blood was obtained prior to and approximately every 30 minutes, when congruent with surgical workflow, during infusion until tissue was removed from the patient. Whole blood was chilled on ice and centrifuged to separate and freeze the plasma. Aliquots of 25-50 μL of plasma were added to 80:20 methanol:water for extraction. Frozen tissue fragments weighing roughly 10-30mg were added to 80:20 methanol:water and extracted to analyze ^13^C enrichment. Samples were subjected to three freeze-thaw cycles, then centrifuged at 16,000xg for 20 minutes to precipitate macromolecules. The supernatant was evaporated using a vacuum concentrator and resuspended in 30 μl of methoxyamine (10 mg/ml) in pyridine. Samples were transferred to autoinjector vials and heated at 70°C for 15 min. A total of 70 μl of tert-butyldimethylsilyl was added, and the samples were briefly vortexed and heated for another 60 min at 70°C. Injections of 1 μl were analyzed on an Agilent 7890A gas chromatograph coupled to an Agilent 5975C mass selective detector. The observed distributions of mass isotopologues were corrected for natural abundance.

### mtDNA: nDNA quantitative polymerase chain reaction (qPCR)

Genomic DNA was isolated using the Qiagen DNeasy Blood & Tissue Kit. Samples were run using the Luna Universal One-Step RT-qPCR Kit (New England Biolabs) on a CFX384 (Bio-Rad). The following primers were used for human COX2 as representative mtDNA: CCGTCTGAACTATCCTGCCC (Forward), (GCCGTAGTCGGTGTACTCGT (Reverse). The following primers were used for human Histone 3 (H4C3) as representative nDNA: GGGATAACATCCAGGGCATT (Forward), CCCTGACGTTTTAGGGCATA (Reverse).

### RNA Isolation

RNA was isolated using Trizol (Thermo Fisher Scientific, Cat. No. 15596018) and an RNeasy Mini Kit (Qiagen, Cat. No. 74106). Total RNA was quantified using a Qubit fluorometer and the Invitrogen Qubit RNA High Sensitivity kit (Invitrogen, Cat. No. Q32852). Samples were diluted in ultrapure water prior to sequencing.

### RNA Sequencing

RNA-seq libraries were prepared using the NEBNext Ultra II directional RNA library prep kit with the NEBNext Poly(A) mRNA magnetic isolation module (New England Biolabs, Cat. No. E7490L, E7760L) according to manufacturer’s instructions. Libraries were stranded using standard N.E.B indices according to manufacturer’s instructions (New England Biolabs, Cat. No. E7730L, E7335L, E7500L). Sequencing reads were aligned to the human reference genome (*hg19*) by STAR 2.7.3.a with default parameters in the 2-pass mod. Counts for each gene were generated using htseq-count v0.6.1. DEGs were identified by DESeq2 v1.14.1. Ends of sequences were trimmed with remaining adapter or quality scores <25. Sequence less than 35bp after trimming were removed. The trimmed Fastq files were aligned to the GRCh38 using HiSAT2^63^ and duplicates were marked with SAMBAMBA. Features (genes, transcripts and exons) were counted using featureCounts^64^. Differential expression analysis was performed using EdgeR^65^ and DESeq^66^. Processed sequencing files will be deposited on GEO. Extended Data Fig. 1A compares RNA sequencing data from the TCGA cohort and this study, emphasizing genes related to the ETC and glycolysis. The Cohen’s effect size (d) between tumour and adjacent kidney for each of 15,642 genes was correlated between the TCGA data and data from the current cohort. ETC genes were selected from the gene ontology cellular component library, including genes related to Complexes I-IV of the ETC. The glycolysis genes include the following four gene sets: KEGG_GLYCOLYSIS_GLUCONEOGENESIS; REACTOME_GLYCOLYSIS; HALLMARK_GLYCOLYSIS; and WP_GLYCOLYSIS_AND_GLUCONEOGENESIS.

### Organotypic Slice Cultures

After surgery, kidney cortex and tumour fragments were embedded in 0.1% agarose and sliced into ∼300 μM thick sections using a microtome (Precisionary Instruments, Copresstome, VF-300). These tissues were then transferred and maintained on hydrophilic PTFE cell culture inserts in human plasma like medium (HPLM) supplemented with 10% dialyzed human serum. Prior to tracing assays, tissues were washed twice with 0.9% saline and medium was replaced with HPLM containing [U-^13^C]glucose for 3 hours. Slices were maintained in an incubator with 5% CO2, 5% O2, and 90% N2.

### Human Serum Dialysis

Human serum was purchased from Sigma-Aldrich (Cat. No. H3667) and dialyzed using SnakeSkin dialysis tubing, 3.5K MWCO, 35 mm (Thermo Fisher Scientific, Cat. No. PI88244). Serum was dialyzed against a 20X volume of PBS. Dialysis was performed for 48 hr at 4°C with a complete PBS exchange every 9-12 hr. Dialyzed serum was then sterile filtered using bottle-top vacuum filters with a pore size of 0.22 μm (Corning Cat. No. 431097).

### Mitochondrial Isolation and Respiration Measurements

Oxygen consumption rates (OCR) were measured using a Seahorse XFe96 Analyzer (Agilent Technologies) as previously described^36,67^. Fresh kidney and tumour samples were homogenized with 40 strokes of a Dounce homogenizer in mitochondrial isolation buffer (HEPES [5 mM], sucrose [70 mM], mannitol [220 mM], MgCl2 [5 mM], KH2PO4 [10 mM], and EGTA [1 mM], pH 7.2) and isolated via differential centrifugation at 4 °C. Nuclei and cell debris were removed by centrifuging five times at 600 xg. Mitochondria were pelleted with a 10000 xg spin and washed twice. 5μg of mitochondria were plated in an XFe96 plate on ice and centrifuged at 2700 xg for 2 minutes at 4 °C. Media containing ETC complex substrates (below) were added to cells and measurements started immediately. At the times indicated, ADP (final concentration 4mM), oligomycin (2 μM), CCCP (2 μM), and either antimycin A (4 μM) or sodium azide (40 μM) were injected. Respiratory control ratios (RCR) were calculated by State III/State IV respiration.

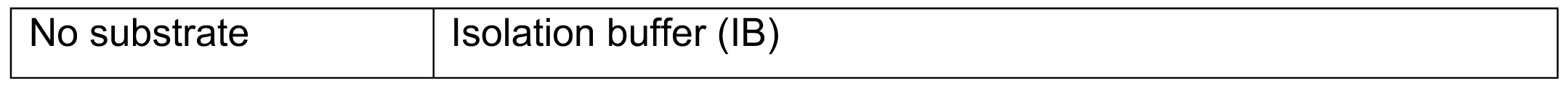
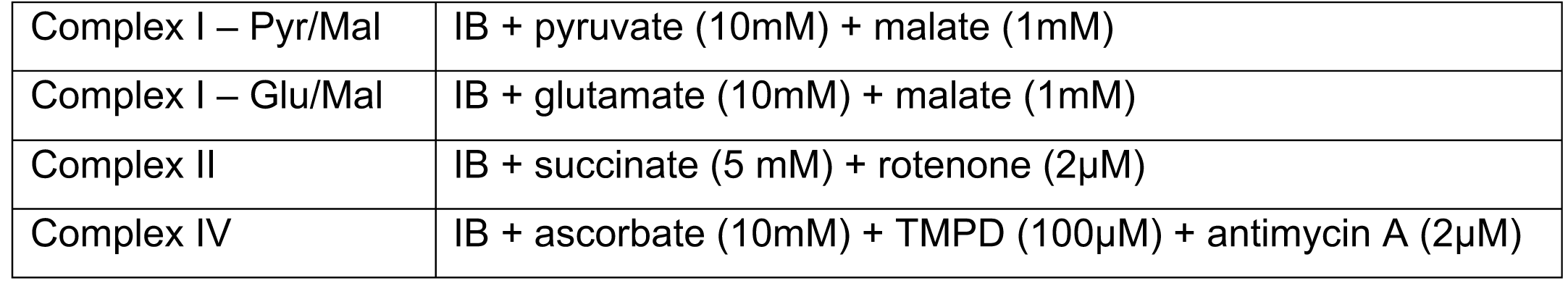

|  |  |
| --- | --- |
| No substrate | Isolation buffer (IB) |
| Complex I – Pyr/Mal | IB + pyruvate (10mM) + malate (1mM) |
| Complex I – Glu/Mal | IB + glutamate (10mM) + malate (1mM) |
| Complex II | IB + succinate (5 mM) + rotenone (2μM) |
| Complex IV | IB + ascorbate (10mM) + TMPD (100μM) + antimycin A (2μM) |

For 786-O cells expressing either empty vector or NDI1, cells were plated in a 96 well plate at a concentration of 2×10^4^ cells/well in 80 μL RPMI-1640 media with 4mM glutamine and 10% FBS. Cells were incubated in a CO2-free incubator at 37°C for 1 hour prior to XFe96 measurements to allow for temperature and pH equilibration. XF assays consisted of 3 mix (3 min) and measurement (3 min) cycles, allowing for determination of OCR/ECAR every 6 minutes.

### NDI1 and dsRed Luciferase Expression

PMXS-NDI1 was a gift from David Sabatini (AddGene, plasmid #72876)^68^. PMXS-NDI1 or PMXS empty vector with gag-pol and VSVG were transfected into 293FT cells using Lipofectamine 3000 (Thermo Fisher, Cat. No. L3000015). Viral supernatants were collected 48 hours after transfection and filtered through a 0.45-μm filter. 786-O cells were cultured with virus containing media and 4µg/mL polybrene (Sigma Aldrich, Cat. No. TR-1003-G) for 24 hours, after which media was changed to fresh media. Cells were then exposed to 10µg/mL blasticidin selection until uninfected 786-O cells died.

A bi-cistronic lentiviral construct carrying dsRed2 and luciferase (dsRed2-P2A-Luc) was a gift from Sean J. Morrison’s laboratory. dsRed2-P2A-Luc with pMD2G and psPAX2 were transfected into 293FT cells using Polyjet (Signagen Cat. No. SL100688) according to manufacturer’s instructions. Viral supernatants were collected 48 hours after transfection and filtered through a 0.45-μm filter. 786-O cells with either PMXS-NDI1 or PMXS empty vector were cultured with virus containing media and 4µg/mL polybrene for 8 hours, after which media was changed to fresh media.

### [U-^13^C]glucose Tracing in Cell Lines

[U-^13^C]glucose tracing data from non-small cell lung cancer cell lines were previously reported^49^. Similar assay conditions were used for tracing experiments in this study. Prior to tracing experiments, 786-O cells expressing either empty vector or NDI1 were washed twice with 0.9% saline and medium was replaced with RPMI-1640 containing [U-^13^C]glucose supplemented with 5% dialyzed FBS for 6 hours. Cells were rinsed in ice cold 0.9% saline and lysed with three freeze thaw cycles in cold 80% methanol. Samples were then prepared for GC/MS analysis.

### Metastatic colonization experiments in mice

All mouse experiments complied with all relevant ethical regulations and were performed according to protocols approved by the Institutional Animal Care and Use Committee at the University of Texas Southwestern Medical Center (Protocol 2016-101360). Cell suspensions were prepared for injection in staining medium (L15 medium containing bovine serum albumin (1 mg/ml), 1% penicillin/streptomycin and 10 mM HEPES (pH 7.4). Tail vein injections were performed in NOD.CB17-*Prkdc^scid^ Il2rg^tm1Wjl^*/SzJ (NSG) mice in a final volume of 50 μL. Four-to-eight-week-old male and female NSG mice were transplanted with 250,000 cells. Both male and female mice were used. Metastatic burden was assessed weekly by bioluminescence. Five minutes before performing luminescence imaging, mice were injected intraperitoneally with 100 μL of PBS containing d-luciferin monopotassium salt (40 mg ml−1; Biosynth, L8220) and mice were anaesthetized with isoflurane 2 min before imaging. The mice were imaged using an IVIS Imaging System 200 Series (Caliper Life Sciences). The exposure time ranged from 10 to 60 s, depending on the maximum signal intensity, to avoid saturation. The bioluminescence signal (total photon flux) was quantified with ‘region of interest’ measurement tools in Living Image software (Perkin Elmer).

### Statistical Analysis

Samples were analyzed as described in the figure legends. Data were considered significant if p<0.05. Statistics were calculated using PRISM software, and statistical details can be found in the figure legends for each figure.

## Extended Data Figure Legends

**Extended Data Figure 1:**
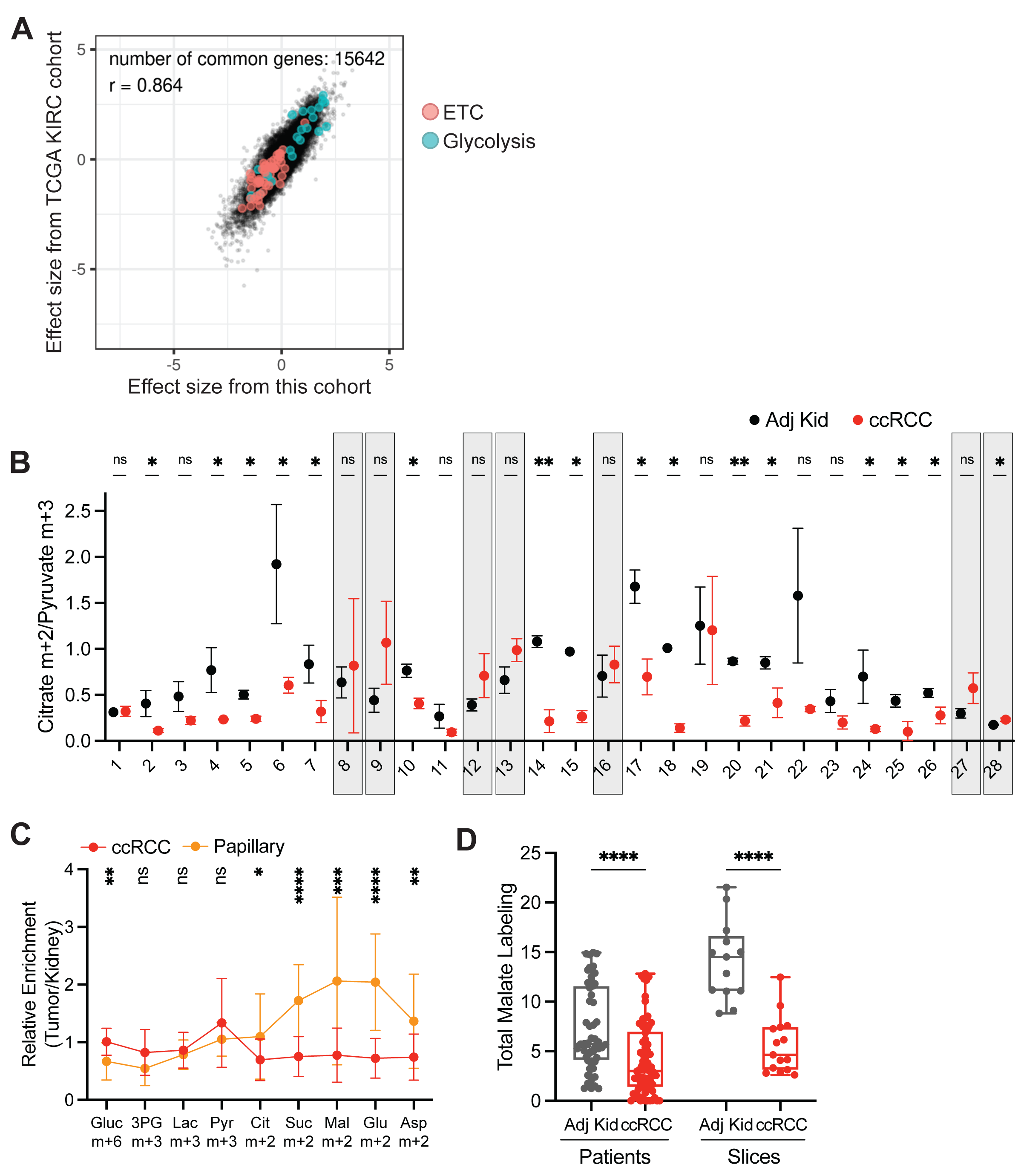
Studied ccRCC tumours reflect heterogenous ccRCC biology. **(A)** Correlation of RNA sequencing data from the TCGA KIRC cohort reporting ccRCC tumours versus the ccRCC tumours infused with [U-^13^C]glucose. Data are plotted as the effect size (Cohen’s d) reflecting the increase (d>0) or decrease (d<0) in mRNA abundance in tumours relative to adjacent kidney. Genes involved in glycolysis and the electron transport chain (ETC) are highlighted as indicated. **(B)** Matched citrate m+2/pyruvate m+3 ratio from patients infused with [U-^13^C]glucose. The x-axis indicates 28 different patients in whom both tumour and kidney tissue was available. Patients in whom the average citrate m+2/Pyruvate m+3 ratio was higher in ccRCC tissue are highlighted in grey boxes; this difference reached statistical significance only in patient 28. **(C)** Enrichment in glycolytic and TCA cycle intermediates associated with glucose oxidation for ccRCC and papillary tumours. Labelling is normalized to the matched adjacent kidney. **(D)** Total malate labeling (1-(m+0)) from [U-^13^C]glucose in patients or tissue slices after 3 hours of labeling. All data represent mean ± standard deviation. Statistical significance was assessed using unpaired t-tests (A-C). \**P* < 0.05, \*\**P* < 0.01, \*\*\**P* < 0.001, \*\*\*\**P*<0.0001. Adj Kid = adjacent kidney, ccRCC = clear cell renal cell carcinoma

**Extended Data Figure 2:**
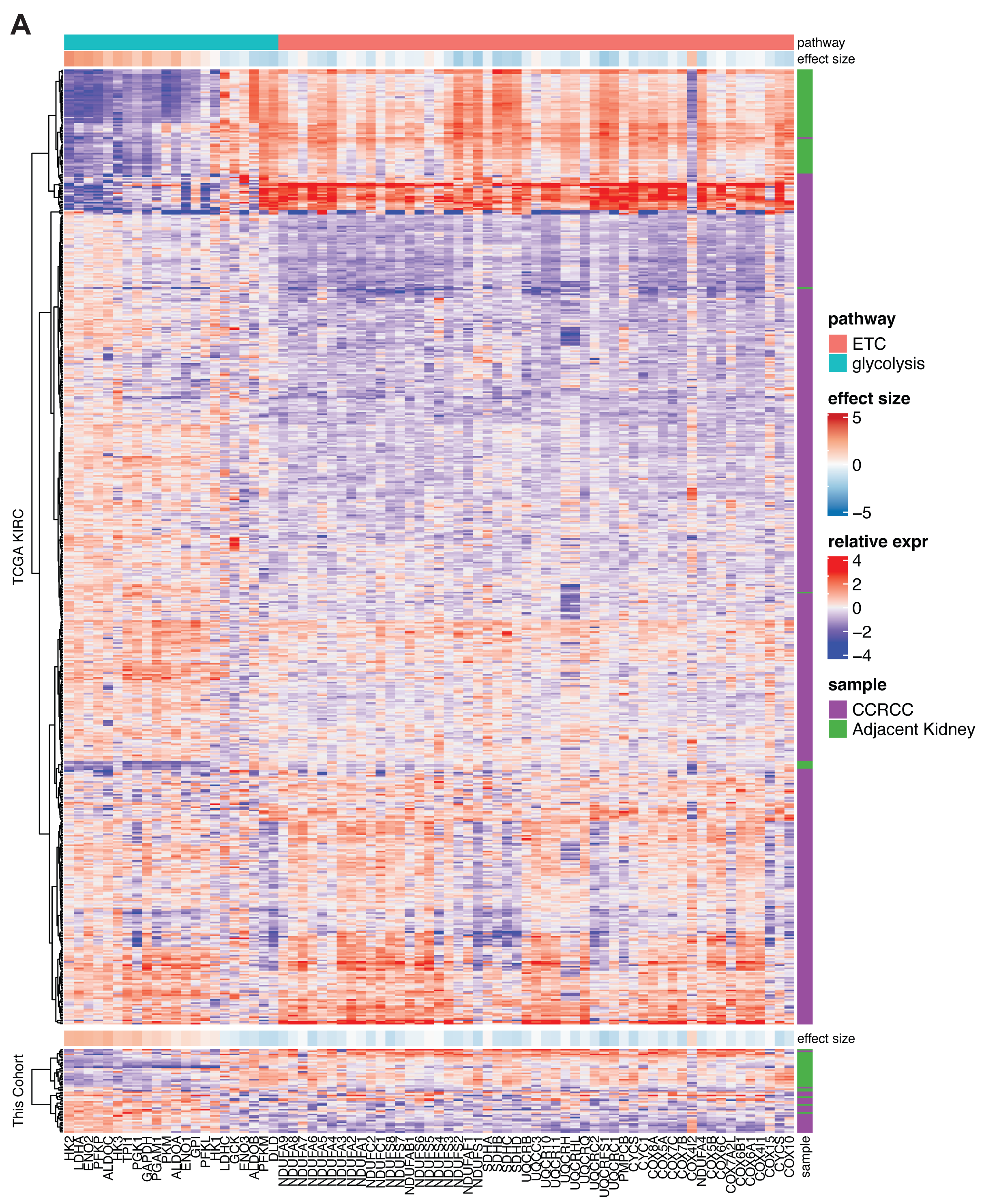
mRNA abundance of ETC and glycolysis genes in primary ccRCC tumours. **(A)** mRNA abundance for genes related to glycolysis and the electron transport chain (ETC) in the TCGA KIRC cohort versus the cohort infused with [U-^13^C]glucose in this study. The ETC genes were selected from the gene ontology cellular component (cc) library combining Complex I-IV. The glycolysis genes are shared genes among the following four gene sets: KEGG_GLYCOLYSIS_GLUCONEOGENESIS, REACTOME_GLYCOLYSIS, HALLMARK_GLYCOLYSIS, WP_GLYCOLYSIS_AND_GLUCONEOGENESIS.

**Extended Data Figure 3:**
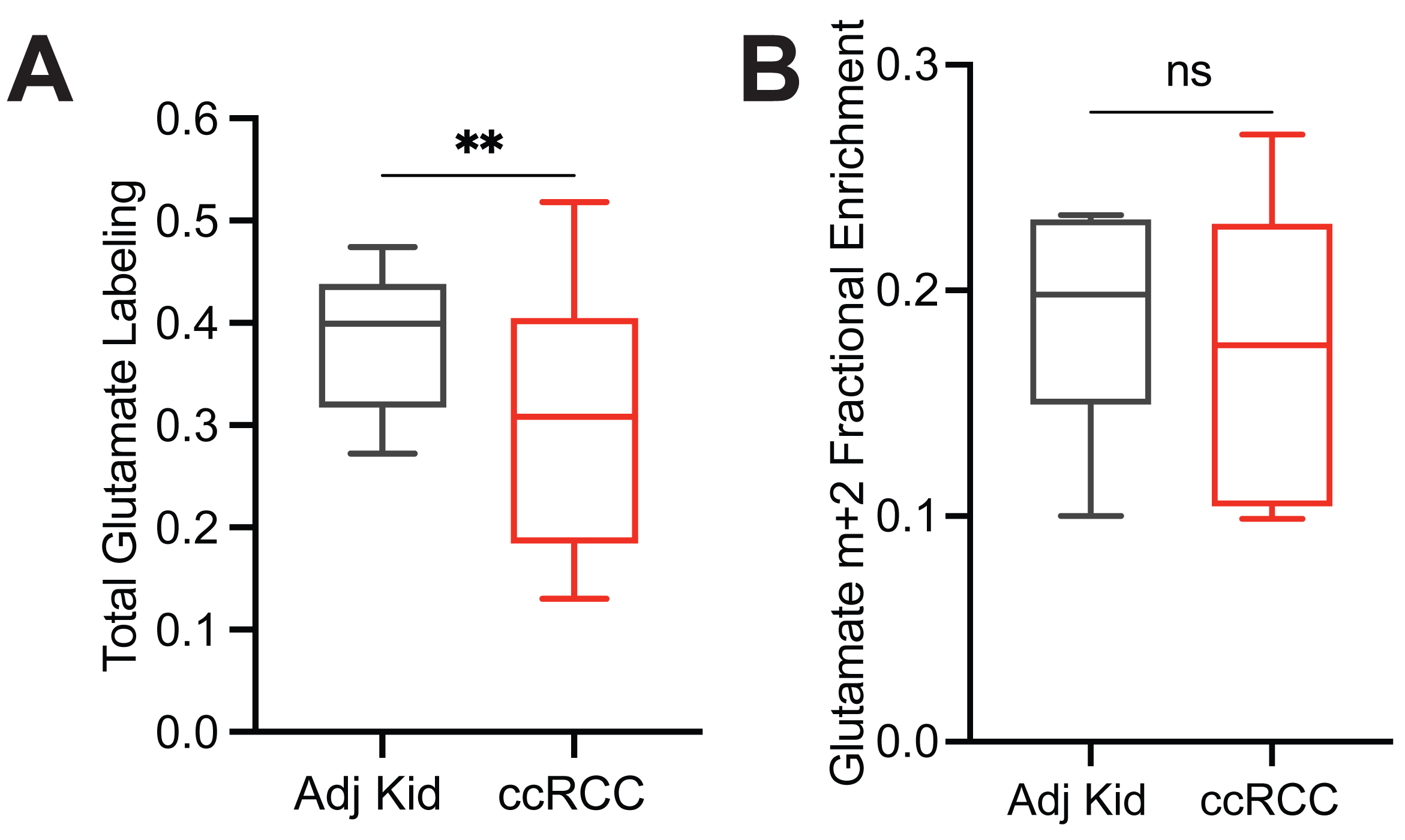
Glutamate enrichment in acetate infused patients. **(A)** Total labeling (1-(m+0)) of glutamate from ccRCC patients infused with [1,2-^13^C]acetate. **(B)** Fractional abundance of glutamate m+2 from ccRCC patients infused with [1,2-^13^C]acetate. Statistical significance was assessed using unpaired t-tests (A, B). \**P* <0.05, \*\**P* < 0.01, \*\*\**P* < 0.001, \*\*\*\**P*<0.0001. Adj Kid = adjacent kidney, ccRCC = clear cell renal cell carcinoma

**Extended Data Figure 4:**
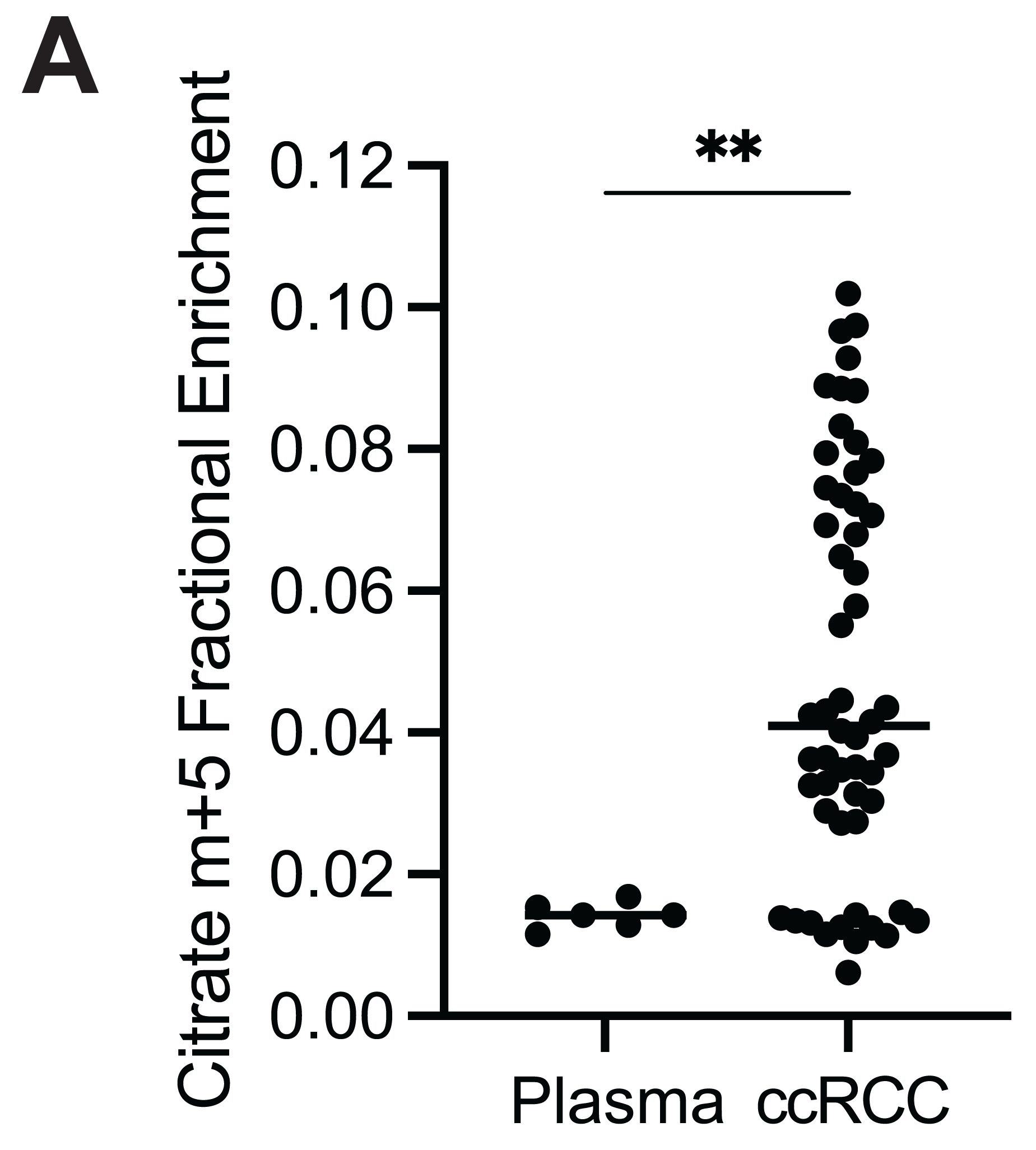
Citrate enrichment in plasma of glutamine infused patients. **(A)** Fractional abundance of citrate m+5 in plasma at the time of resection and in ccRCC tumour samples. Statistical significance was assessed using unpaired t-tests (A, B). \**P* < 0.05, \*\**P* < 0.01, \*\*\**P* < 0.001, \*\*\*\**P*<0.0001. ccRCC = clear cell renal cell carcinoma

**Extended Data Figure 5:**
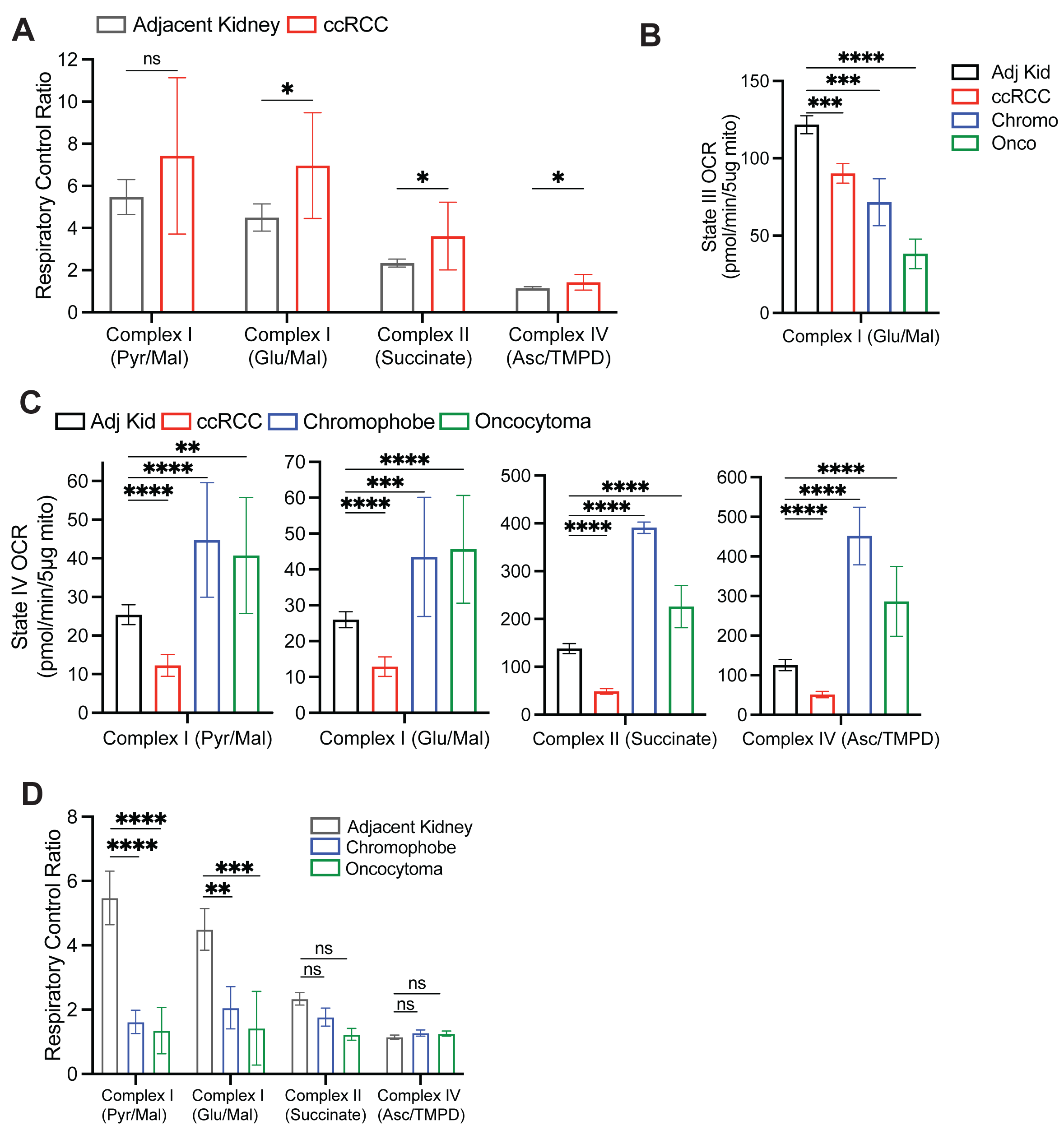
Respiration of primary human kidney cancers. **(A)** Respiratory control ratio (RCR) for mitochondria from the adjacent kidney and ccRCCs. RCR is the ratio of State III ADP-stimulated OCR to the State IV basal OCR. **(B)** State III ADP-stimulated oxygen consumption rates (OCR) from mitochondria isolated from primary human tissues, using glutamate and malate to stimulate Complex I. **(C)** State IV basal OCR from mitochondria isolated from primary human tissues. Injected substrates are indicated under each complex. **(D)** Respiratory control ratio (RCR) for chromophobe RCCs and oncocytomas. Panels A-C represent mean ± 95% confidence intervals, and panel D represents mean ± standard deviation. Statistical significance was assessed using an unpaired two tailed parametric t-test (A) or one way analysis of variance (ANOVA) with a multiple comparison adjustment using Tukey’s methods (B-D). ns *P>*0.05, \**P* < 0.05, \*\**P* < 0.01, \*\*\**P* < 0.001, \*\*\*\**P*<0.0001. TMPD = N,N,N,N-tetramethyl-p-phenylenediamine.

**Extended Data Figure 6:**
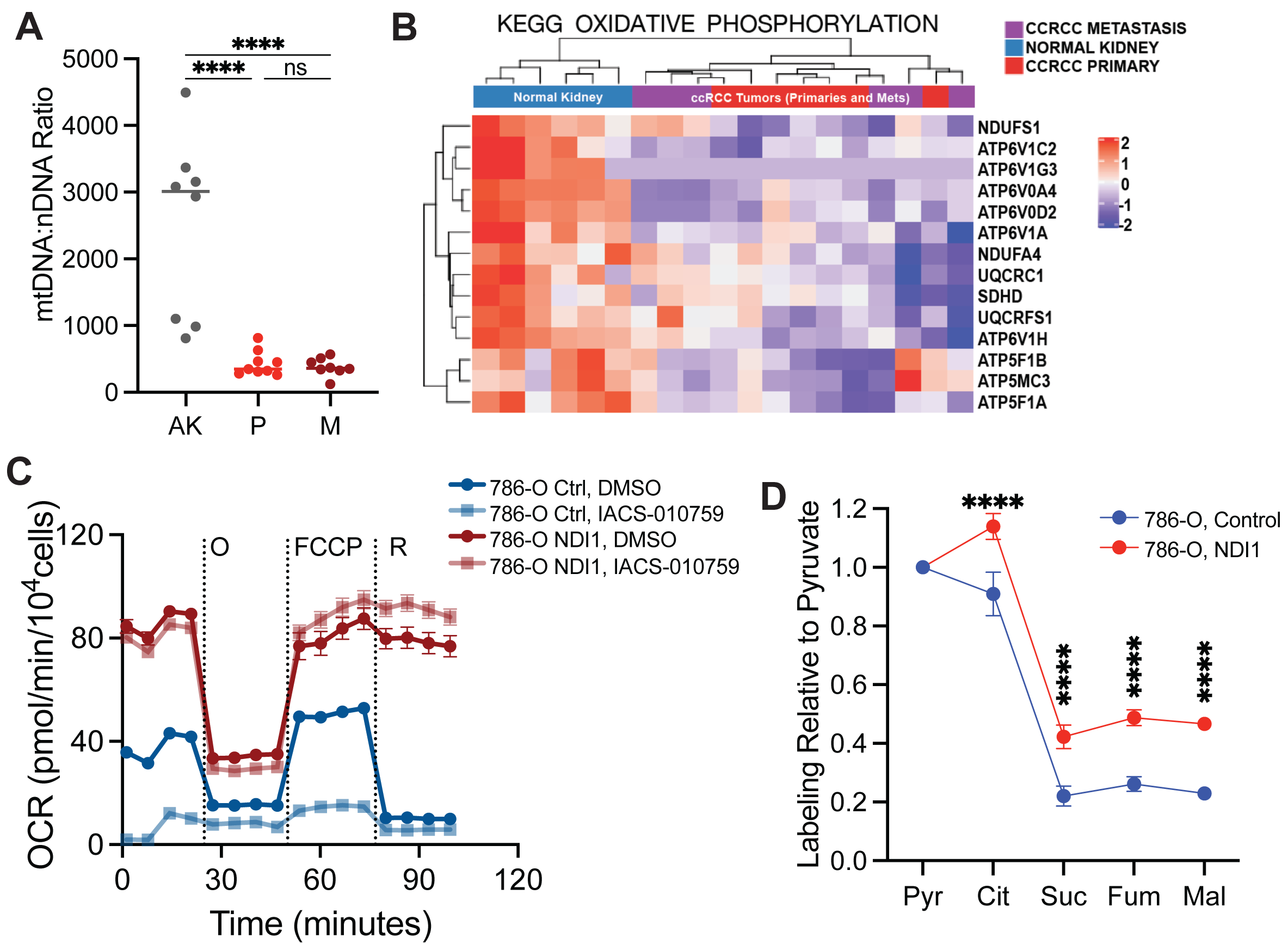
Mitochondrial characteristics in metastasizing ccRCC tumours and ccRCC cells. **(A)** mtDNA:nDNA ratio from 7 patients from the adjacent kidney (AK), primary ccRCC (P), and metastastic ccRCC (M). **(B)** Heat map of the most differentially expressed genes in the oxidative phosphorylation gene set from RNA sequencing of the 7 matched patients in Extended Data Fig 6A. **(C)** Oxygen consumption rates of 786-O cells expressing either the control empty vector or NDI-1. IACS-010759 is a Complex I inhibitor. **(D)** Total labeling in TCA cycle intermediates relative to pyruvate. Statistical significance was assessed using an unpaired two tailed parametric t-test (A) or one way analysis of variance (ANOVA) with a multiple comparison adjustment using Tukey’s methods (B-D). ns *P>*0.05, \**P* < 0.05, \*\**P* < 0.01, \*\*\**P* < 0.001, \*\*\*\**P*<0.0001. O = oligomycin, FCCP = carbonyl cyanide-p-trifluoromethoxyphenyl-hydrazon, R = rotenone.

